# Ontology based mining of pathogen-disease associations from literature

**DOI:** 10.1101/437558

**Authors:** Șenay Kafkas, Robert Hoehndorf

## Abstract

**Background:** Infectious diseases claim millions of lives especially in the developing countries each year, and resistance to drugs is an emerging threat worldwide. Identification of causative pathogens accurately and rapidly plays a key role in the success of treatment. To support infectious disease research and mechanisms of infection, there is a need for an open resource on pathogen-disease associations that can be utilized in computational studies. A large number of pathogen-disease associations is available from the literature in unstructured form and we need automated methods to extract the data.

**Results:** We developed a text mining system designed for extracting pathogen-disease relations from literature. Our approach utilizes background knowledge from an ontology and statistical methods for extracting associations between pathogens and diseases. In total, we extracted a total of 3,420 pathogen-disease associations from literature. We integrated our literature-derived associations into a database which links pathogens to their phenotypes for supporting infectious disease research.

**Conclusions:** To the best of our knowledge, we present the first study focusing on extracting pathogen-disease associations from publications. We believe the text mined data can be utilized as a valuable resource for infectious disease research. All the data is publicly available from https://github.com/bio-ontology-research-group/padimi and through a public SPARQL endpoint from http://patho.phenomebrowser.net/.

## Background

Each year, millions of people die due to infectious diseases. The World Health Organisation (WHO)[1] reported that 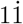 million deaths were due to HIV/AIDS in 2015 alone. Infectious diseases cause devastating results not only on global public health but also on the countries economies. Developing countries, especially the ones in Africa, are the most affected by infectious diseases.

Several scientific resources have been developed to support infectious disease research. A large number of these resources focus on host–pathogen interactions [2, 3] as well as particular mechanisms of drug resistance [4]. Additionally, there are several resources that broadly characterize different aspects of diseases [5]. Relatively little structured information is available about the relationships between pathogens and disease, information that is also needed to support infectious disease research. Currently, such associations are mainly covered by proprietary databases such as the Kyoto Encyclopedia of Genes and Genomes (KEGG) [6] as well as the biomedical literature and public resources such as Wikipedia [7], MedScape [8], or the Human Disease Ontology [5] in natural language form. Automated methods are needed to extract the associations from natural language.

Here, we further developed and evaluated a text mining system for extracting pathogen–disease associations from literature[9]. While most of the existing text mining studies related to infectious disease focus on extracting host–pathogen interactions from text [10, 11] and archiving this data [2, 3], to the best of our knowledge, we present the first text mining system which focuses on extracting pathogen-5 disease associations. Our literature-extracted associations are available for download from https://github.com/bio-ontology-research-group/padimi and through a public SPARQL endpoint at http://patho.phenomebrowser.net/.

## Materials & Methods

### Ontologies and Resources Used

We used the latest archived version of the Open Access full text articles (http://europepmc.org/ftp/archive/v.2017.12/, containing approximately 1.8 million articles) from the Europe PMC database [12]. We used the NCBI Taxonomy [13] (downloaded on 22-08-2017) and the Human Disease Ontology (DO) [5] (February 2018 release) to provide the vocabulary to identify pathogen and infectious disease mentions in text. We generated two dictionaries from the labels and synonyms in the two ontologies and refined them before applying text mining. In the refinement process, we filtered out terms which have less than three characters and terms that are ambiguous with common English words (e.g., “Arabia” as a pathogen name). We extracted only the species labels and synonyms belonging to fungi, virus, bacteria, worms, insects, and protozoa from NCBI Taxonomy to form our pathogen dictionary. The final pathogen and disease dictionaries cover a total of 1,250,373 distinct pathogens and 438 distinct infectious diseases.

### Pathogen and Disease class Recognition

A class is an entity in an ontology that characterizes a category of things with particular characteristics. Classes usually have a set of terms attached as labels or synonyms [14]. We used the Whatizit text mining pipeline [15] to annotate pathogen and disease classes in text with the two dictionaries for diseases and pathogens. Because disease name abbreviations can be ambiguous with some other names (e.g., ALS is an abbreviation both for “Amyotrophic Lateral Sclerosis” and “Advanced Life Support”), we used a disease abbreviation filter for screening out the ambiguous abbreviations that could be introduced during the annotation process [16]. Briefly, this filter operates based on rules utilizing heuristic information. First, it identifies abbreviations and their long forms in text by using regular expressions. Second, it utilizes several rules to decide whether to keep the abbreviation annotated as a disease name or filter out. The rules cover keeping the abbreviation either if any of its long forms from DO exists in the document or its long form contains a keyword such as “disease”, “disorder”, “syndrome”, “defect”, etcṫhat describes a disease name.

### Pathogen–Disease Association Extraction

Our association extraction method is based on identification of pathogen-disease co-occurrences at the sentence level and applying a filter based on co-occurrence statistics and Normalized Point-wise Mutual Information (NPMI) [17] association strength measurement to reduce noise possibly introduced by the high recall, low precision co-occurrence method. We selected the associations having an NMPI value above 0.2 and co–occurring at least 10 times in the literature.

We extended NPMI, which is a measure of collocation between two terms, to a measure of collocation between two classes. Hence, we reformulated the NPMI 8 measure for our application. First, we identify, for every class, the set of labels and synonyms associated with the class (*Labels*(*C*) denotes the set of labels and synonyms of *C*). We then define *Terms*(*C*) as the set of all terms that can be used to refer to *C*: *Terms*(*C*) := {*x*|*x* ∈ *Labels*(*S*) ∧ *S* ⊑ *C*}.

We calculate the NPMI between classes *C* and *D* as

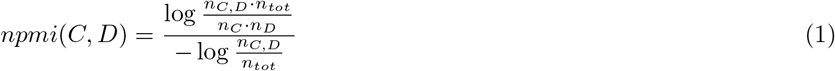

where *n*_*tot*_ is the total number of sentences in our corpus (i.e., 4,427,138), *n*_*C,D*_ is the number of sentences in which both a term from *Terms*(*C*) and a term from *T erms*(*D*) co-occur, *n*_*C*_ is the number of sentences in which a term from *Terms*(*C*) occurs, and *n*_*D*_ is the number of sentences in which a term from *Terms*(*D*) occurs.

## Results

### Statistics on Extracted Pathogen–Disease Associations

We extracted a total of 3,420 distinct pathogen–disease pairs from over 1.8 million Open Access full text articles. To identify the associations, we used a combination of lexical, statistical, and ontology-based rules. We used lexical matches to identify whether the label or synonym of a pathogen or disease is mentioned in a document; we used a statistical measure, the normalized point-wise mutual information, to determine whether pathogen and disease mentions co–occur significantly often in literature; and we used ontologies as background knowledge to expand sets of terms based on ontology-base inheritance.

### Performance Evaluation

To evaluate the pathogen–disease associations we obtain, we used the KEGG [6] database as reference and compare our results to the information contained in KEGG. KEGG contains pathogen–disease associations obtained through manual curation. We could identify 744 pathogen–disease pairs in KEGG for which we could map the pathogen and disease identifiers from NCBI Taxonomy and DO to their identifiers in KEGG. Figure 1 shows the overlapping and distinctly identified pathogen-disease associations from Kegg and literature. We covered 29.4% (219) of the pathogen–disease associations from KEGG and extracted many more associations from literature (3,201). There are 525 pairs which we could not cover by text mining. The main reason we cannot identify an association is due to limitations in our named entity and normalization procedure.

**Figure 1.**
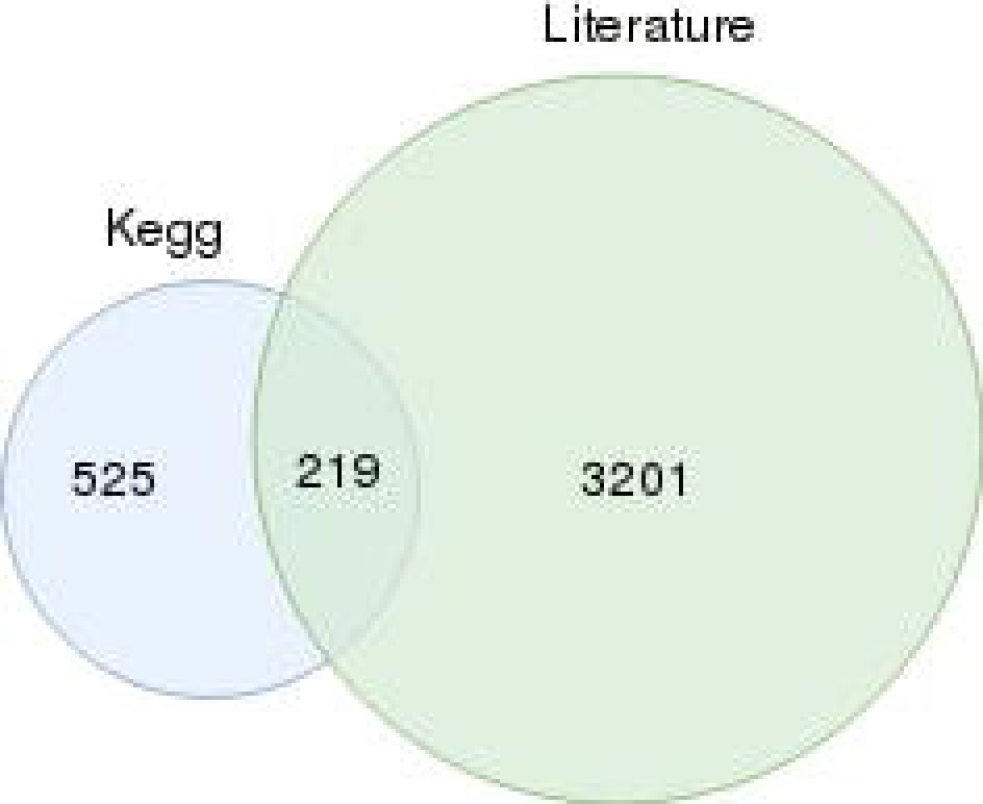
Overlapping pathogen–disease associations between Kegg and literature

We further evaluated the performance of our system manually on 50 randomly selected pathogen–disease associations. In our manual evaluation, we achieve a precision of 64%, a recall of 84% and an F-score of 73%. The false positives were mainly due to ambiguous abbreviations and pathogen names. For example, “bronchus” was annotated as an insect name by our method.

Some false negatives were due to rejections by the pipeline based on the threshold settings. For example, “Tetranychus” and “asthma” co-occurred only once in our corpus and therefore the association between them was rejected as we limited our analysis to pathogen–disease pairs that co-occurred ten or more times. Other false negatives were due to missing pathogen or disease labels in our dictionaries.

## Discussion

By using ontologies as background knowledge to expand our sets of terms and labels, it is possible to identify pathogen–disease associations even if the labels and synonyms directly associated with the pathogen or disease are not directly found to co–occur in text. For example, we extracted a total of 44 distinct pathogen–disease associations relevant to *dengue disease* (`DOID:11205`). 12/44 of these associations are the direct associations of *dengue disease* (i.e., a label or synonym of the disease is explicitly mentioned in text) while the remaining 32/44 are indirect associations obtained from associations with labels and synonyms of the sub-classes *asymptomatic dengue* (`DOID:0050143`), *dengue hemorrhagic fever* (`DOID:12206`), and *dengue shock syndrome* (`:0050125`). In total, we found 812 pathogen-disease associations which do not directly co-occur in literature.

The performance of our system depends on two parameters: the NPMI value and the number of co-occurrences used as a threshold. In the future, we may use these two values to automatically determine optimal threshold based on a more comprehensive evaluation set of pathogen–disease associations. While our initial text mining approach performs at a promising level (F-score 73%), there is still some room for improvements. As we found the pathogen names to be ambiguous with other domain specific names, we plan to further improve the abbreviation and name filters we apply. For improving the recall of our system, it may be possible to expand our dictionaries with other resources covering disease and pathogen names such as the Experimental Factor Ontology (EFO) [18] and the Unified Medical 132 Language System (UMLS) [19] for diseases, and the Encyclopedia of Life [20] for 133 pathogens.

## Conclusion

Here, we present a text mining method for extracting pathogen-disease associations from the biomedical literature. Our method performed at the state of the art level with some room for improvements. In future, we plan to improve our text mining method by developing and integrating a pathogen abbreviation filter and expanding the coverage of our pathogen and disease dictionaries. In the scope of infectious disease research, we have included our results in a database of pathogens and the phenotypes they elicit in humans. We believe that our results can further support infectious disease research.

## List of abbreviations

(WHO): World Health Organisation
(KEGG): Kyoto Encyclopedia of Genes and Genomes
(DO): Human Disease Ontology
(NPMI): Normalized Point-wise Mutual Information
(EFO): Experimental Factor Ontology
(UMLS): Unified Medical Language System

## Availability

All the data is available from https://github.com/bio-ontology-research-group/padimi and (http://patho.phenomebrowser.net/) through a public SPARQL endpoint.

## Author’s contributions

RH and SȘK conceived of the study; SȘK performed all experiments. ȘK and RH analyzed the results. ȘK drafted the manuscript, R.H. revised the manuscript. All authors have read and approved the final version of the manuscript.

## Consent of publication

Not applicable.

## Ethics approval and consent to participate

Not applicable.

## Acknowledgement

Authors would like to thank Mrs. Marwa Abdellatif for her help to make the data available from the SPARQL end-point.

## References

1. World Health Organisation. http://who.int/en/

2. Ammari, M.G., Gresham, C.R., McCarthy, F.M., Nanduri, B.: HPIDB 2.0: a curated database for host-pathogen interactions. Database 2016(2016)

3. Wardehand, M., Risley, C., McIntyre, M.K., Setzkorn, C., Baylis, M.: Database of host-pathogen and related species interactions, and their global distribution. Scientific Data 2(150049, eCollection2015) (2015)

4. Jia, B., Raphenya, A.R., Alcock, B., Waglechner, N., Guo, P., Tsang, K.K., Lago, B.A., Dave, B.M., Pereira, S., Sharma, A.N., Doshi, S., Courtot, M., Lo, R., Williams, L.E., Frye, J.G., Elsayegh, T., Sardar, D., Westman, E.L., Pawlowski, A.C., Johnson, T.A., Brinkman, F.S.L., Wright, G.D., McArthur, A.G.: Card 2017: expansion and model-centric curation of the comprehensive antibiotic resistance database. Nucleic Acids Research 45(D1), 566–573 (2017)

5. Kibbe, W.A., Arze, C., Felix, V., Mitraka, E., Bolton, E., Fu, G., Mungall, C.J., Binder, J.X., Malone, J., Vasant, D., Parkinson, H.E., Schriml, L.M.: Disease ontology 2015 update: an expanded and updated database of human diseases for linking biomedical knowledge through disease data. Nucleic Acids Research 43(Database-Issue), 1071–1078 (2015)

6. Kanehisa, M., Goto, S.: KEGG: kyoto encyclopedia of genes and genomes. Nucleic Acids Research 28(1), 27–30 (2000)

7. List of Infectious Diseases. https://en.wikipedia.org/wiki/List_of_infectious_diseases

8. Medscape. https://emedicine.medscape.com/

9. Kafkas, S., Hoehndorf, R.: Ontology based mining of pathogen – disease associations from literature. In: Hoenhdorf, R., Dumontier, M. (eds.) Proceedings of Bio-Ontologies SIG@ISMB 2018, 6-10 July 2018; Chicago, USA. (2018)

10. Thieu, T., Joshi, S., Warren, S., Korkin, D.: Literature mining of host-pathogen interactions: comparing feature-based supervised learning and language-based approaches. Bioinformatics 28(6), 867–875 (2012)

11. İlknur Karadeniz, Hur, J., He, Y., Özgür, A.: Literature mining and ontology based analysis of host-brucella gene-gene interaction network. Frontiers in Microbiology 6(1386) (2015)

12. Europe PMC: a full-text literature database for the life sciences and platform for innovation. Nucleic Acids Research 43(Database-Issue), 1042–1048 (2015)

13. Sayers, E.W., Barrett, T., Benson, D.A., Bryant, S.H., Canese, K., Chetvernin, V., Church, D.M., DiCuccio, M., Edgar, R., Federhen, S., Feolo, M., Geer, L.Y., Helmberg, W., Kapustin, Y., Landsman, D., Lipman, D.J., Madden, T.L., Maglott, D.R., Miller, V., Karsch-Mizrachi, I., Ostell, J., Pruitt, K.D., Schuler, G.D., Sequeira, E., Sherry, S.T., Shumway, M., Sirotkin, K., Souvorov, A., Starchenko, G., Tatusova, T.A., Wagner, L., Yaschenko, E., Ye, J.: Database resources of the national center for biotechnology information. Nucleic Acids Research 37(Database-Issue), 5–15 (2009)

14. Hoehndorf, R., Schofield, P.N., Gkoutos, G.V.: The role of ontologies in biological and biomedical research: a functional perspective. Briefings in Bioinformatics 16(6), 1069–1080 (2015)

15. Rebholz-Schuhmann, D., Arregui, M., Gaudan, S., Kirsch, H., Jimeno, A.: Text processing through web services: calling whatizit. Bioinformatics 24(2), 296–298 (2008)

16. Kafkas, S., Dunham, I., McEntyre, J.R.: Literature evidence in open targets – a target validation platform. J. Biomedical Semantics 8(1), 20–1209 (2017)

17. Bouma, G.: Normalized (pointwise) mutual information in collocation extraction. In: Proceedings of the Biennial GSCL Conference: 2009; Potsdam, Germany pp. 31–40 (2009)

18. Malone, J., Holloway, E., Adamusiak, T., Kapushesky, M., Zheng, J., Kolesnikov, N., Zhukova, A., Brazma, A., Parkinson, H.E.: Modeling sample variables with an experimental factor ontology. Bioinformatics 26(8), 1112–1118 (2010)

19. Bodenreider, O.: The unified medical language system (UMLS): integrating biomedical terminology. Nucleic Acids Research 32(Database-Issue), 267–270 (2004)

20. Encylopedia of Life. http://eol.org/

